# GATLncLoc+C&S: Prediction of LncRNA subcellular localization based on corrective graph attention network

**DOI:** 10.1101/2024.03.08.584063

**Authors:** Xi Deng, Lin Tang, Lin Liu

## Abstract

Long non-coding RNAs (LncRNAs) have a wide range of regulatory roles in gene expression, and the subcellular localization identification of LncRNAs is of great value in understanding their biological functions. Graph neural networks can not only utilize sequence characteristics, but also learn hidden features from non-Euclidean data structures to obtain features with powerful characterization capabilities. To learn more fully from the limited LncRNA localization samples and efficiently exploit easily ignored label features, we propose a corrective graph attention network prediction model GATLncLoc+C&S in this paper. Compared with previous methods, the similarity of optimal features is first used to construct the graph. Then, a re-weighted graph attention network R-GAT is constructed and the soft labels obtained from it are used to correct the graph. Finally, the predicted localization label is further obtained by label propagation. Based on the combination of R-GAT and label propagation, GATLncLoc+C&S effectively solves the problems of few samples and data imbalance in LncRNA subcellular localization. The accuracy of GATLncLoc+C&S reached 95.8% and 96.8% in the experiments of 5- and 4-localization benchmark datasets, which reflects the great potential of our proposed method in predicting LncRNA subcellular localization. The source code and data of GATLncLoc+C&S are available at https://github.com/GATLncLoc-C-S/GATLncLoc-C-S.

## 1 Introduction

Long non-coding RNA (LncRNA) is a non-coding RNA more significant than 200 nucleotides in length [1]. Human lncRNAs participate in a spectrum of biological processes, for example, epigenetics, nuclear import, alternative splicing, RNA decay, and translation [2,3]. They can also serve as precursors to small RNAs. Therefore, aberrant lncRNA expression can cause various human diseases and disorders [4]. Increasing reports of dysregulated LncRNA expression in many cancer types imply that LncRNAs may act as potential tumor suppressor RNAs [3,5]. Many studies have shown that LncRNAs are highly adaptable molecules capable of biological functions through RNA-RNA, RNA-DNA, or RNA-protein interactions. Several LncRNAs localized in the nucleus or chromatin have been experimentally shown to regulate the transcriptional expression of adjacent genes in cis, or in trans to regulate the transcription of a subclass of genes through interactions between heterogeneous ribonucleic acid proteins (hnRNPs) [6,7,8,9]. Furthermore, several LncRNAs in the cytoplasm have been shown to interact with RNAs and proteins to perform their molecular functions [6,10]. For example, Cesana et al. [11] found that LincMD1 is mainly expressed in the cytoplasm of differentiated muscle cells and acts as a competitive RNA (ceRNA) to regulate the differentiation process of skeletal muscle. It can be seen that LncRNAs are present in a variety of subcellular structures of cells, and specific subcellular localization is essential for the biological functions of LncRNAs. Since relying on biochemical experiments to determine lncRNA subcellular localization information is time-consuming and expensive, it is imperative to use efficient and stable computational models to predict lncRNA subcellular locations.

Currently, few studies are based on computational models to predict the subcellular localization of LncRNAs. From a methodological point of view, it is more based on traditional machine learning. On the one hand, traditional machine learning-based methods focus more on feature extraction aspects. For example, reference [16] extracted sequence features based on the 8-mer and PseKNC [22] methods and used a binomial distribution-based approach to reduce the feature dimensionality by feature selection to build the optimal feature subset; reference[18] was based on k-mer, PseDNC [23] and triplet methods to extract sequence features, and then the combination of the above three features is used for feature selection using methods such as variance thresholding and binomial distribution; Reference[15] extracted LncRNA sequence features based on k-mer while learning a high-level representation of the original parts using stacked autoencoders. The experimental results showed that the extracted deep features can discover hidden feature correlations from the original high-dimensional features with better discriminative power. On the other hand, current traditional machine learning-based methods mostly utilized random forests, support vector machines (SVMs), and logistic regression to accomplish classification tasks. In the 4 classification task, references [16][17] were based on SVM to complete the classification prediction, where to solve the sample imbalance problem, reference [17] obtained a high accuracy of 90.69% by two SMOTEs. Reference [15] obtained 66.5% accuracy by random forest, support vector machine (SVM) and self-encoder using only one sequence feature extraction method; Reference[18] used three sequence feature extraction methods based on logistic regression to achieve high accuracy of 92.37%. It can be seen that the fusion based on multiple sequence features can establish the association between markers and location labels more accurately and comprehensively than the traditional classifiers, and the in-depth exploration of the extraction and selection of sequence features can further assist the traditional classifiers to obtain superior prediction performance. Meanwhile, with the impressive performance of deep learning methods in automatically capturing advanced data features, several studies on predicting LncRNA subcellular localization have also adopted deep learning methods in recent years. For example, reference [24] adopted multilayer deep neural networks for both nucleus and cytoplasm localization; reference [19] proposed an integrated model IDDLncLoc based on convolutional neural networks and SVM; reference [20] obtained a complete feature representation through the embedding of subsequences and applied a text convolutional neural network (textCNN) to learn high-level features. Thus, obtaining more discriminative classification features based on the robust feature fusion and mining ability of deep learning is the primary development trend in this field.

With the development of LncRNA subcellular localization studies, several related databases have emerged, including RNALocate [12], LncATLAS [13], and LncSLdb [14]. These databases provide a solid foundation for studying computational models of LncRNA subcellular localization. Among them, the RNAlocate database contains more subcellular locations of LncRNAs and is the source of data for most LncRNA subcellular localization prediction experiments. There is 9587 LncRNA localization information in the RNAlocate database, of which 6636 species belong to human LncRNAs. However, by counting these databases, we found that the distribution of LncRNA subcellular localization data showed an extremely unbalanced trend (as shown in Fig.1). In particular, the number of LncRNA samples under each localization label is more drastically reduced after screening LncRNA samples based on data preprocessing. For example, the dataset in reference [15-20] has just over 600 entries after preprocessing and is in 4 or 5 localization labels. Therefore, even though deep learning can automatically capture high-level features but cannot achieve better results in lncRNA subcellular localization prediction tasks due to the small number of training samples and imbalance problem. For example, reference [20] obtained an accuracy of 53.7% in a 5classification task. Currently, for this problem, reference [15] [17] [19] used a synthetic minority class oversampling algorithm (SMOTE) to alleviate the training difficulties of the model, and reference [19] reached a high accuracy of 94.96%. However, while the model’s predictive capability is improved, the SMOTE algorithm will have a particular impact on the distribution of the original data, with problems such as overfitting. Also, for a small amount of data, there needs to be more data to represent the class boundaries, and the final expanded decision region remains error-prone by SMOTE synthesis of noise and boundary examples [49]. It can be seen that the lack of training samples and data imbalance problems are more significant in deep learning, which remains a serious problem for LncRNA subcellular location prediction at this stage.

**Fig. 1.**
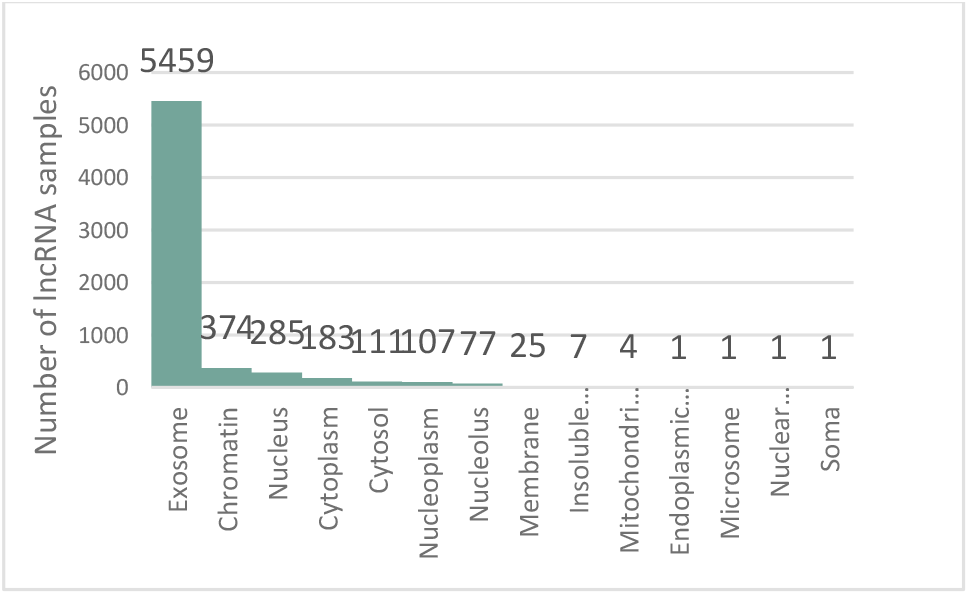
Statistics of LncRNA subcellular location data.

**Fig. 2.**
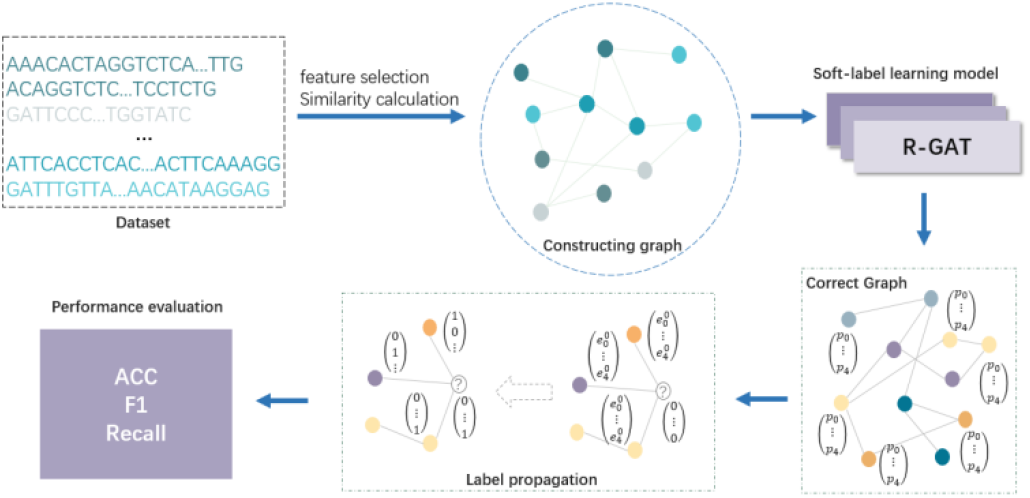
The flow chart of GATLncLoc+C&S.

Unlike the general two-dimensional matrix feature, the graph structure has a more robust representation capability. It can represent a variety of rich and complex data and is considered a structured data type characterized by vertices (entities that hold information) and edges (connections between vertices that contain information) with compositional and relational properties. In recent years, combining graph neural networks has led to extraordinary achievements in disease prediction and drug research[25].

Inspired by this, we propose a new idea to predict lncRNA subcellular localization using graph neural networks and label propagation. The LncRNA sequence features are converted into graph structures, and on the one hand, the LncRNA sequence features and spatial features are combined into components with strong expression ability by graph neural networks. On the other hand, due to the costly acquisition of labeled data in biological experiments, label propagation (LPA) based on the idea that neighboring sample points have the same label is convenient and can effectively improve the predictive power of models. LPA is currently achieving encouraging results in many fields, such as multimedia information retrieval, classification, and online community discovery [26, 27]. In this paper, the optimal features are first obtained from the sequence features by recursive feature elimination, and the adjust cosine similarity of the optimal features is calculated to construct the initial graph structure. Since each node has different contributions from neighboring nodes, the graph attention network GAT [28] can be used to learn node embeddings by introducing an attention mechanism to estimate the importance coefficient between each node and the nodes associated with it. For the data imbalance problem, we try to combine the class weights calculated based on the Effective Number of Samples proposed by Cui et al. [29] and the Label-Distribution-Aware Margin Loss function (LDAM) [30] in GAT model, so as to construct a soft label learning model R-GAT, which provides larger margins for the minority classes by better understanding the feature expressions of the minority classes. In addition, the sequence conservation of LncRNA is low, and their sequence similarity is similar to the intron region of protein-coding genes, which is below 70% in humans and mice and slightly lower than the 5′ or 3′ untranslated regions of genes [31]. As a result, feature similarity-based conformation can exist where fewer nodes with the same localization label between neighborhood nodes, leading to insufficient feature learning of nodes by R-GAT, which can be misleading for label propagation based on the network having homogeneity or coordination. This paper proposes a soft label corrective graph structure obtained using RGAT to address this problem. Meanwhile, inspired by the label correction and smoothing C&S methods proposed by Huang [32] et al., this paper obtains the final prediction results based on the corrective graph structure by propagating the label errors in the training set to correct the error information in the test set and then smoothing the prediction on the test set.

Based on the above ideas, GATLncLoc+C&S in this study mainly includes the following modules: (1) constructing the benchmark dataset; (2) constructing the initial graph structure; (3) soft label learning model R-GAT; (4) correcting the graph structure; (5) label propagation; and (6) performance evaluation.

## 2 Materials and Methods

### 2.1 Benchmark Dataset

RNALocate (http://www.rna-society.org/rnalocate/) [12] records many RNA-related subcellular localization entries and provides a user-friendly interface for users to query and browse detailed information on RNA subcellular localization. With the starting point of providing more details on LncRNA localization, we used the 5-classified dataset Dataset1 constructed by Cao et al. [15], containing 612 sequences of LncRNAs. To reduce information redundancy and noise interference, one series of length 91671 and 11 lines containing special symbols “N, R, S, Y” were deleted to obtain the dataset Dataset2, which includes 600 LncRNAs, including 292 Cytoplasm, 149 Nucleus, 91 Cytosol, 43 Ribosome, and 25 Exosome sequences, as shown in Table 1.

**Table 1.**
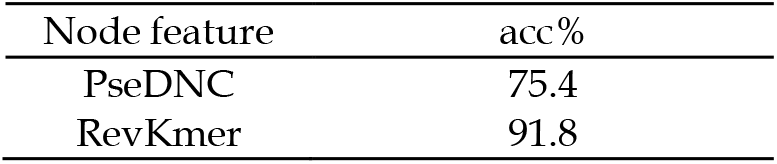
Benchmark dataset.

The common sequence feature extraction methods currently used in LncRNA subcellular localization prediction include k-mer [33], RevKmer [34,35], PseDNC [36-38], Triplet [39-41], and DACC [19]. After the experimental comparison, the specific experimental results are detailed in Section 3.1; we finally chose k-mer to represent the features. K-mer can represent local sequence order information of DNA or RNA sequences and is a commonly used and effective feature extraction method in various fields of bioinformatics [42-45], which yields a high-dimensional feature vector of dimension 4^k^ by calculating the frequency of occurrence of k-tuple nucleotides in a sequence [46-48].

When we set k to 6, 7, and 8, the noise or redundancy present in this high-dimensional data will affect the model accuracy and generalization ability, so the representative low-dimensional feature vector is obtained by recursive feature elimination.

### 2.2 Construction of the initial graph

The representation of graph structure is obtained from the optimal feature vector of LncRNA sequences, which enables the model to get more hidden information from non-Euclidean space, and the graph structure with high homogeneity is conducive to better learning of the model. The graph consists of three parts: the set of nodes V, the set of node features X, and the set of edges E. The initial graph is defined as G = (V, E, X), Node set V = {v_1_, v_2_, …, v_m_} ; Edge Set E = {e_1,2_, e_1,3_, …, e_I,j_} ; Feature set X = {x_1_, x_2_, x_3_, …, x_I_}, x_I_ denotes the optimal feature vector of node v_I_ in the initial graph G; The set of labels Y ∈ {1,2, …, C} indicates that there are class C subcellular locations. Our goal is to predict the subcellular location y_i_ ∈ *Y* of a lncRNA (v_i_ ∈ *V*) by aggregating information about the node features (x_i_) of the lncRNA (v_i_) and the features of its neighboring nodes.

#### 2.2.1 Calculating the optimal set of edges

Due to the low similarity of LncRNA sequences, we choose the Adjust Cosine Similarity, which considers both vector directions and is more sensitive to the value, and obtain the similarity matrix S by decentering and then calculating the cosine similarity. M denotes the total number of samples in this experiment. Finally, the symmetric matrix of S(_m*m)_ is obtained, S(_i*m)_ or S(_m*I)_ indicates the similarity between the i-th node and the rest of the nodes, and the n highest similarities S(_i*m)_ or S(_m*I)_ are selected from S(_i*j)_, to determine the edge that creates the i-th node and the j-th node (e_I,j_ = 1; otherwise, e_I,j_ = 0), as in Equation (1).

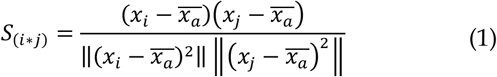

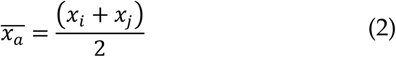

#### 2.2.2 Visualization of the initial graph

Based on the edge set determination of the initial graph above, we used five colors to represent the five localizations of cytoplasm, nucleus, cytosol, ribosome, and exosome, respectively, and the extreme imbalance of the data can also be seen more intuitively in Fig.3.

**Fig. 3.**
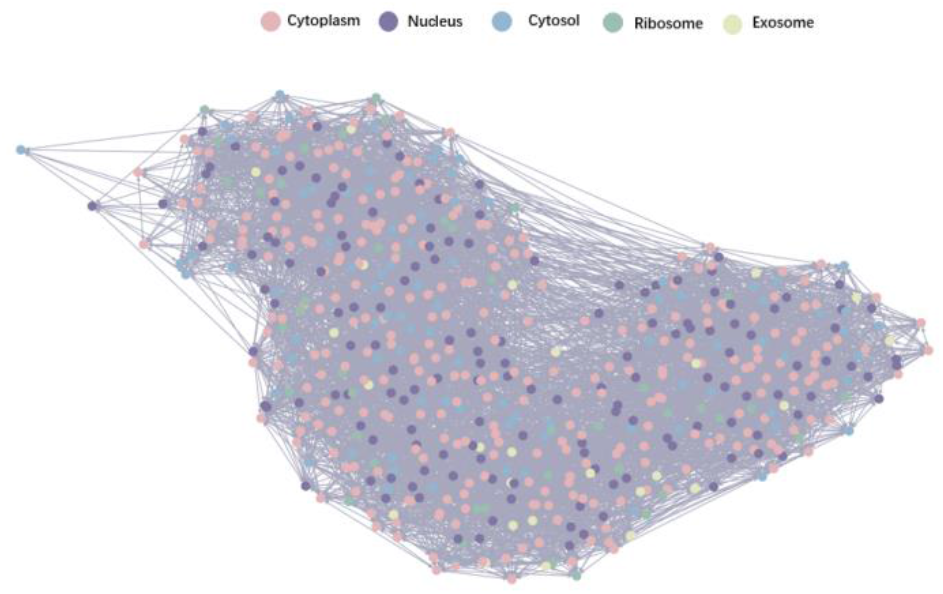
Visualization of the initial diagram.

### 2.3 Soft label learning model R-GAT

The LncRNA sequence features are transformed into graph structures. Then the graph neural network is used to aggregate the information of the graph containing topological relationships to obtain more expressive new features, including the original feature information and connection information. In this paper, we propose the soft label learning model R-GAT, which sets weights for the information aggregation between nodes and their neighboring nodes through an attention mechanism, thus making a hierarchical distinction between the interconnections of nodes. Combined with the newly introduced re-weighting method, the model can better learn the information of fewer classes of nodes thus improving the prediction accuracy.

Each node v_I_ in the model corresponds to a LncRNA, and the optimal features are selected from the k-mer features. The initial graph structure is constructed by the calculation of similarity, with the input initial graph G = (V, E, X) and the set of node feature vectors.

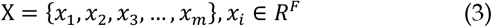

F is the optimal feature dimension after dimensionality reduction, and the feature set X^′^ of the new dimension is output by attention coefficient calculation and feature aggregation:

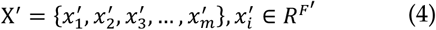

The attention coefficient θ_ij_ after softmax normalization is calculated as follows:

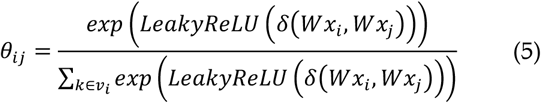

In the above Equation, k ∈ v_I_ means k is an adjacent node of node vI, the weight matrix W ∈ R^F*F′^ implements a linear transformation from F dimensional input features to F^′^ dimensional output features, δ is a learnable weight vector, and LeakyReLU is a nonlinear-activation-function. The new features 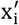 for each node are output from feature aggregation as follows:

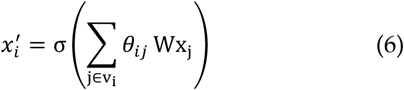

In this paper, a multi-headed attention mechanism is used to average the K-independent attention mechanisms to make the self-attentive process more stable, which acts as an integration to prevent overfitting and enables to enrich the feature extraction capability of the model compared to a single head:

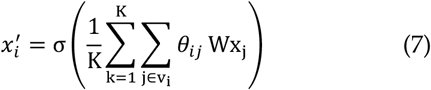

To alleviate the data imbalance problem, R-GAT utilizes a class balancing weight [29] method with an inversely proportional number of effective samples, associating each piece with a small neighborhood. Meanwhile, R-GAT combines the L_LDAM_ loss [30] to find the optimal balance from the uneven edges of the unbalanced dataset, providing more prominent advantages for a few classes by minimizing the edge-based generalization boundary:

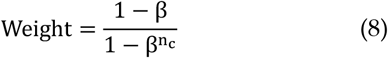

n_c_ denotes the number of samples in class c, the hyperparameter β ∈ (0,1), which is generally set to [0.9, 1) for better results in most experiments; in this experiment, beta =0.999.

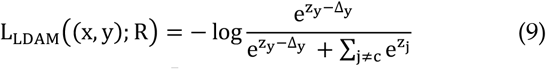

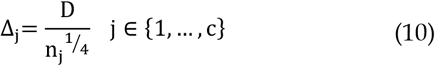

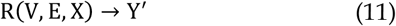

where R is shorthand for the soft label learning model R-GAT, D is the hyperparameter, n_j_ is the number of samples in each class, and z_y_ is the probability distribution of the model output. The parameters are continuously updated by gradient return to improve the learning ability of R-GAT for features and obtain more accurate soft labels Y^′^, preserving the soft label learning model R-GAT with the highest accuracy in the validation process.

### 2.4 Correctional diagram structure

Then the saved soft label learning model R-GAT is loaded, and the similarity matrix S of soft labels Y^′^ between nodes is calculated:

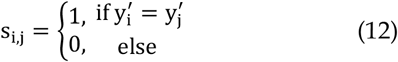

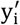 denotes the soft label of node I; if the soft labels y^′^ of nodes v_I_ and v_j_ are the same, then S_I,j_ = 1 ; otherwise, S_I,j_ = 0. The edges of nodes v_I_ and v_j_ are determined by randomly selecting h nodes from S_I,j_ = 1 (e_I,j_ = 1; otherwise, e_I,j_ = 0). P_I_ denotes the set of class probabilities corresponding to 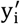.In five classifications, P_i_ = {p_0_, p_1_, p_2_, p_3_, p_4_}; at this time, the feature set of the nodes in the graph is updated to the corresponding set of class probabilities P = {P_1_, P_2_, P_3_, …, P_m_}, obtaining the corrective graph G^′^ = (V, E, P).

### 2.5 Label propagation

We obtain the training error vector e for each node by using the difference between one-hot encoding y_I_ of the true labels in the training set and the class probability P_I_ corresponding to soft labels, and initialize the training error e to 0 for nodes without labels:

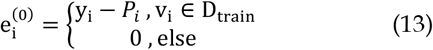

The training error is propagated through the graph G^′^, and the error is iterated t times using the adjacency matrix A, the degree matrix D, and the training error matrix E. The hyperparameter β determines the degree or proportion of information retained from the previous step, and the hyperparameter γ adjusts the balance of the error matrix after the iteration to obtain the corrected soft label Y^′′^:

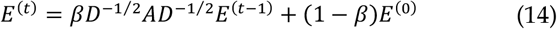

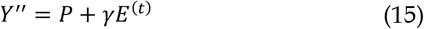

Finally, based on the great possibility that neighboring nodes belong to the same class, we replace the corrected soft label Y^′′^ with the true label Y for the training set nodes to obtain a new label matrix Z. Using the idea of information transfer, the label matrix Z is iterated t times to get the label matrix Z^(t)^, and the one with the highest class probability is selected from this matrix as the final test set for classification:

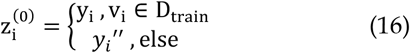

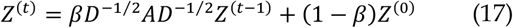

The algorithm flow of the GATLncLoc+C&S model is shown in Fig.4:

**Fig. 4.**
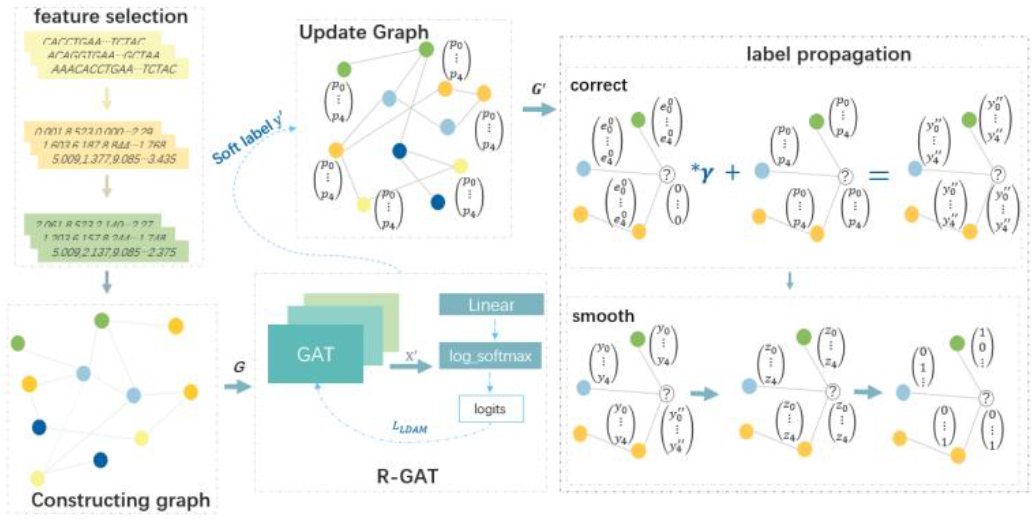
Algorithm flow chart of GATLncLoc+C&S.

### 2.6 Performance evaluation

So far, the standard metrics for evaluating LncRNA subcellular localization models are Accuracy (Acc), Recall (R), F1 score (F1), sensitivity (Sn), specificity (Sp), and Matthews’s correlation coefficient ( MCC) [16,17,18]etc. To better compare with other methods and more objectively evaluate the performance of GATLncLoc+C&S, we assessed the following metrics of GATLncLoc+C&S by a 5-fold cross-validation method:

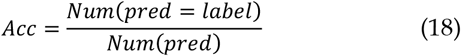

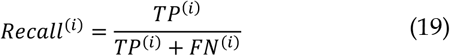

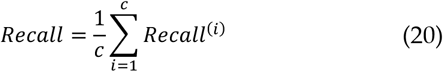

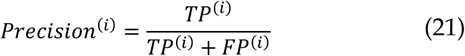

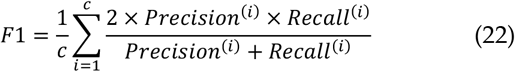

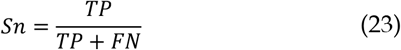

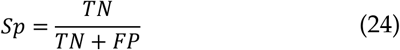

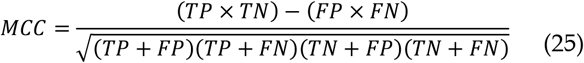

TP^(i)^, TN^(I)^, FP^(I)^, and FN^(I)^ denote the true positives, true negatives, false positives, and false negatives of class i, c represents the label type.

## 3 Results and Discussion

### 3.1 Dataset partitioning

Due to the peculiarities of small and unbalanced data, we do five-fold cross-validation for each localization label separately to avoid affecting the original data distribution. Each label of data is divided into five subsets. The first subset of each class is combined into a test set, and the remaining four subsets form the training set. The second subset of each class is taken to create a new test set, and the remaining four subsets include the new training set. The third and fourth, in turn, are taken to obtain five copies of D_teSt_ and D_tRain_ . Finally, we adopt D_teSt_ as D_vaL_ and satisfy D_teSt_ ∪ D_tRain_ = D, D_teSt_ ∩ D_tRain_ = ∅.

### 3.2 Comparing the effects of different feature extraction methods on model performance

We initially compared the prediction results of four methods for extracting primary features, including DACC, PseDNC [36-38], RevKmer [34,35], and k-mer [33], where both k-mer and RevKmer take k values of 6, and PseDNC takes λ and ω of 150 and 0.3, respectively. First, the initial graph is constructed by calculating the similarity based on the feature vectors obtained by the above four methods, as seen in Fig.5. The initial graph obtained by the RevKmer and k-mer methods has a higher proportion of neighboring nodes belonging to the same localization label, 63.3% and 64.6%. The initial graph obtained by the PseDNC process shows that the rate of neighboring nodes with the same localization label is low at 49.65%. Then, we constructed the initial graph with the initial features extracted by PseDNC, RevKmer, and k-mer methods. We fixed the set of node features as the feature vector extracted by k-mer to obtain the absolute accuracy of 84.6%, 93.3%, and 94.5%.

**Fig. 5.**
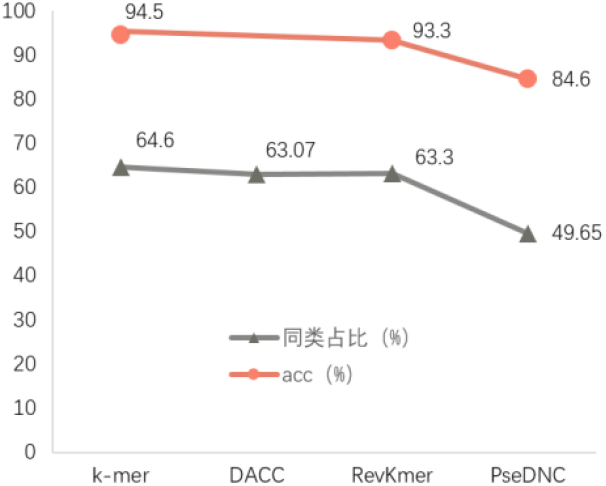
Performance comparison of constructing graph structures based on different feature extraction methods.

From the above experimental results, it is determined that the similarity is better calculated using the feature vectors extracted by k-mer. The initial features extracted with the three methods, PseDNC, RevKmer and k-mer, were compared again as the feature sets of the nodes, and the absolute accuracy was 75.4%, 91.8% and 94.5%, respectively (Table 2). we can deduce that GATLncLoc+C&S has the best performance when both the data for constructing the graph and the node feature set are extracted by k-mer, which is why we choose k-mer as the node feature for prediction.

**Table 2.**
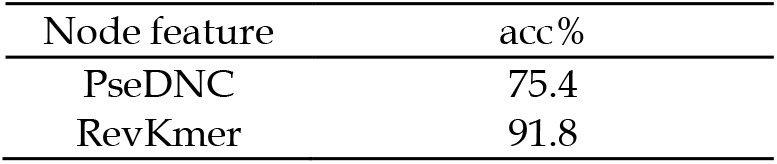
Effect of different feature extraction methods on node feature sets.

### 3.3 Comparing the effect of different numbers of edges in the initial graph G on the model performance

We perform experimental comparisons in three dimensions, feature 6-mer, 7-mer, and 8-mer, and construct different initial graphs based on 20, 30, and 40 neighboring nodes taken for each node n. In this experiment, the percentage of neighboring nodes of the same localization label in the initial graph obtained by taking the different number of edges n per node is first analyzed. It can be seen from Fig.6 that the larger the value of n for each node, the smaller the percentage of neighboring nodes with the same localization label in the initial graph, regardless of the importance of the k-mer taken. When the number of edges n per node is 20, the percentage of neighboring nodes with the same localization label is higher. With the increase of k in k-mer, the rate of neighboring nodes with the same localization label also shows an increasing trend, and when the 8-mer feature is downscaled to 4000, the highest percentage of neighboring nodes with the same localization label is 68.37%. Accordingly, the value of the number of edges n per node affects the rate of neighboring nodes with the same localization label in the graph more than the value of k-mer and the dimensionality of feature selection. Finally, according to the proportion of neighboring nodes with the same localization label in the initial graph in different dimensions obtained from Fig.6, we downscaled the feature 6-mer to 2000 and the feature 7mer and 8-mer dimensions to 4000 and analyzed Acc, Recall, and F1 metrics by GATLncLoc+C&S prediction. The horizontal coordinates in Fig. 7 indicate the combination of different dimensions with the number of edges n taken per node, and it shows that the indicator tends to decrease with increasing n in 6-mer, 7-mer, and 8-mer, and the experiment further confirms that it is better when the number of edges n taken per node is 20. Meanwhile, the highest accuracy of 95.8% was obtained at 8-mer, which confirms that a higher percentage of samples with the same localization label in the neighboring nodes gives a better prediction result. In addition, the number of edges n per node is more important than the feature dimension choice. For example, in 8-mer, as the number of edges per node n is taken from 20 to 40, the difference between the highest accuracy and the lowest accuracy is 5%, while the number of edges per node n is taken as 20, in k-mer as k is taken as 6, 7, and 8, the difference is 2.8%.

**Fig. 6.**
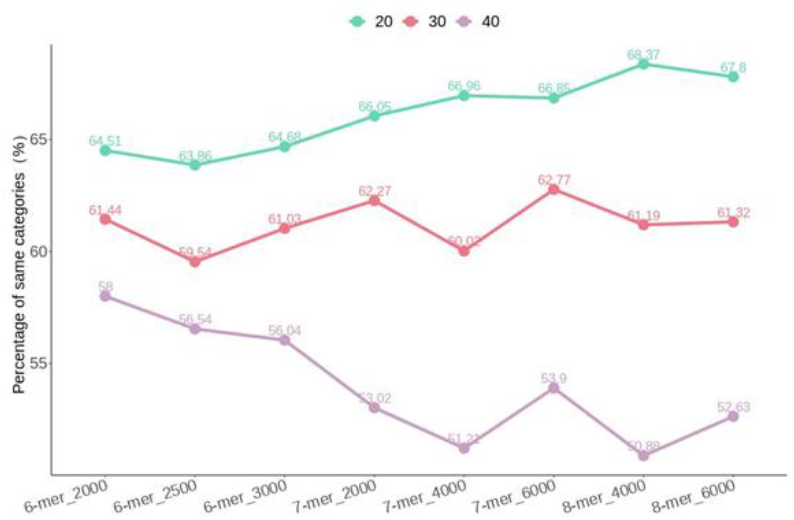
The percentage of nodes of the same in neighboring nodes with different values of the number of edges taken per node.

**Fig. 7.**
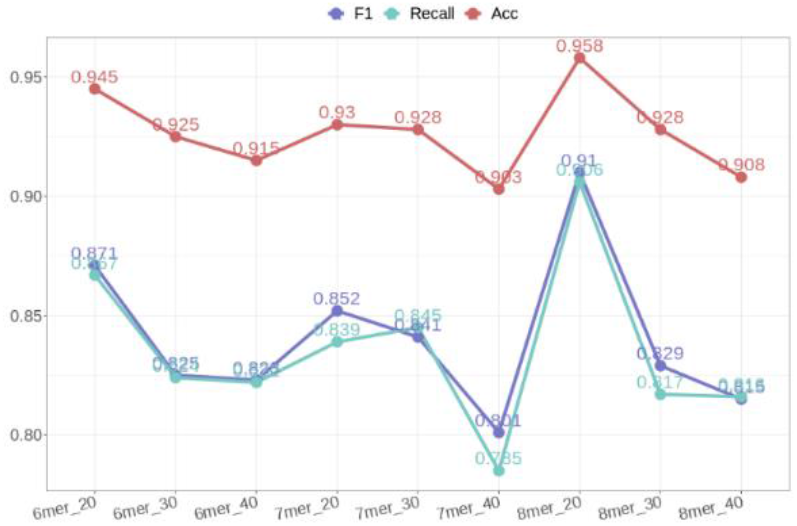
Comparing the effect of different numbers of edges in the initial graph G on the model performance.

### 3.4 Comparing the effect of optimal features of different dimensions on model performance

The best prediction result of taking 20 neighboring nodes for each node is determined by experiment 3.3. The initial graph is constructed by further selecting the optimal features of different dimensions to calculate the similarity, which is found to have less influence on the prediction result of this model. From Table 3, it can be seen that feature 6-mer Acc shows a decreasing trend with a dimensionality increase. For feature 6-mer, the experimental results of dimensionality reduction to 2000 are better, with an accuracy of 94.5%. There is a slight fluctuation of indicators in feature 7-mer; after considering the time cost and equipment conditions, this experiment does not further confirm the specific optimal dimensionality. Feature 8-mer also showed a slight decrease in Acc with the increase of dimensionality, and the evaluation metrics all reached the highest value when the optimal feature was taken 4000, in which the accuracy rate was as high as 95.8%.

**Table 3.** Comparison of model performance of optimal features in different dimensions.

| Dataset with 5 subcellular localization dataset |  |  |  |  |
| --- | --- | --- | --- | --- |
| K-mer | Optimal dimension | F1 | Recall | Acc(%) |
| 6-mer | 2000 | 0.871 | 0.867 | 94.5 |
|  | 2500 | 0.830 | 0.827 | 92.5 |
|  | 3000 | 0.801 | 0.798 | 92.6 |
| 7-mer | 4000 | 0.852 | 0.839 | 93.0 |
|  | 6000 | 0.865 | 0.863 | 94.1 |
|  | 8000 | 0.871 | 0.863 | 93.8 |
| 8-mer | 4000 | **0.910** | **0.906** | **95.8** |
|  | 6000 | 0.887 | 0.870 | 94.9 |
|  | 8000 | 0.875 | 0.868 | 94.6 |

### 3.5 Compare the model performance of R-GAT, R-GAT+C&S, and GATLncLoc+C&S

To confirm the effectiveness of our proposed soft label-based correction of the initial graph structure before label propagation, this experiment compares the prediction results of the soft label learning model R-GAT, label propagation without modification of graph structure R-GAT+C&S and GATLncLoc+C&S models. Based on the optimal parameters selected from the above experiments, this comparison experiment uses feature 8-mer downscaled to 4000 as the experimental data. From Table 4, it is clear that using soft labels to correct the initial graph structure and then doing label propagation has a more significant impact on this experiment. From the soft label learning model R-GAT and the initial graph-based label propagation R-GAT+C&S experiments, it can be seen that the original graph is somewhat misleading for label propagation, and the accuracy drops from the initial 91.3% to 89.6%; the main reason is that there are fewer samples of the same localization label in the neighboring nodes. The three evaluation metrics of the soft label learning models R-GAT and GATLncLoc+C&S show that correcting the initial graph based on soft labels is necessary, and the accuracy rate is improved from the original 91.3% to 95.8% by correcting the initial graph.

**Table 4.**
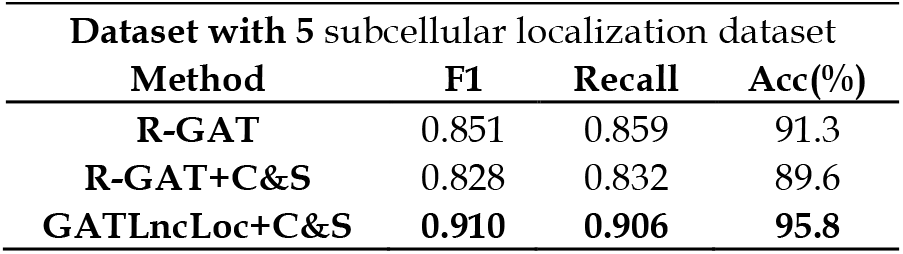
Performance comparison of R-GAT, R-GAT+C&S, and GATLncLoc+C&S.

| Dataset with 5 subcellular localization dataset |  |  |  |
| --- | --- | --- | --- |
| Method | F1 | Recall | Acc(%) |
| R-GAT | 0.851 | 0.859 | 91.3 |
| R-GAT+C&S | 0.828 | 0.832 | 89.6 |
| GATLncLoc+C&S | **0.910** | **0.906** | **95.8** |

### 3.6 Comparing the effect of different numbers of edges in the new graph G’ on the model performance

The comparison of experiments from 3.5 shows that using a soft label correction graph structure significantly affects the subsequent label propagation. Therefore, we further verify the influence of the number of edges per node in graph G^′^ on the existence of the practical effect.

Since the number of samples of the minimum class in this experiment is 25, based on this, we try to set the number of edges h per node as 3, 5, 7, 10, 15, and 20. The horizontal coordinate of Fig.8 is the number of edges h at each node, and it can be seen that all metrics tend to decrease as the number of edges h at each node increases, where all three evaluation metrics reach the highest point when the number of edges at each node in the corrected graph structure is set to 5.

**Fig. 8.**
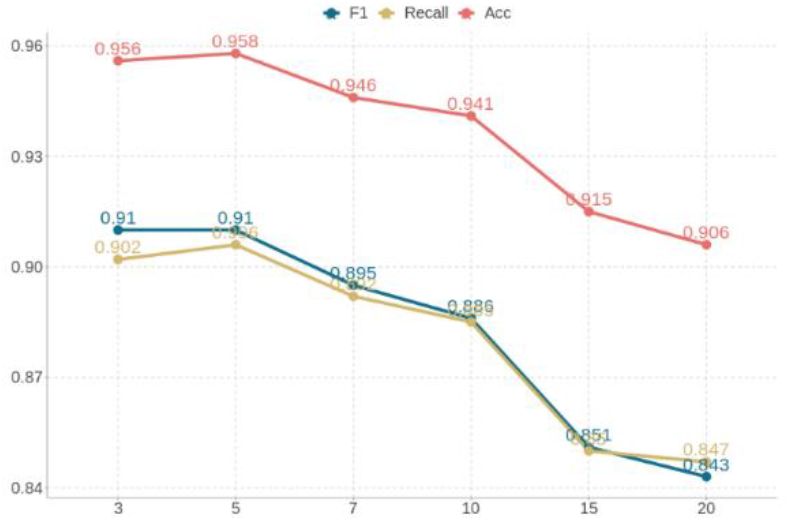
Comparing the effect of different numbers of edges in the new graph G’ on the model performance.

### 3.7 Comparing the performance of other models

The existing models for studying 5-localization (Cytoplasm, Nucleus, Cytosol, Ribosome, and Exosome) are lncLocator [15] and DeepLncLoc [20]. From the comparison of three metrics, F1, Recall, and Acc, as shown in Table 5, the three metrics of GATLncLoc+C&S are much higher than lncLocator and DeepLncLoc. In addition, GATLncLoc+C&S also compared with five predictors on 4-category benchmark datasets (Cytoplasm, Nucleus, Ribosome, and Exosome), including iLoc-lncRNA [16], Locate-R [17], LncLocPred [18], IDDLncLoc [19] and Proposed by Yang et al. [51]. We performed a comparative analysis by four evaluation metrics, Sensitivity, Specificity, MCC, and Acc. It is evident from Table 6 that GATLncLoc+C&S is better compared to the other five predictors in many aspects of evaluation metrics, in which the accuracy of GATLncLoc+C&S is about 1.84% higher than IDDLncLoc, which achieves the best experimental results in comparison models. It further indicates that the proposed GATLncLoc+C&S in this paper has great potential in lncRNA subcellular localization prediction.

**Table 5.**
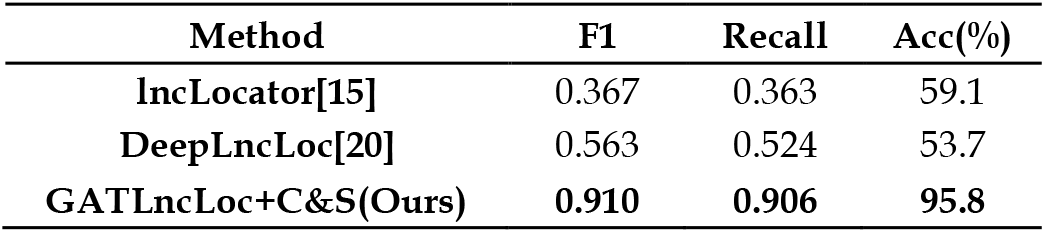
Comparison with state-of-the-art methods on 5 subcellular localization dataset.

**Table 6.**
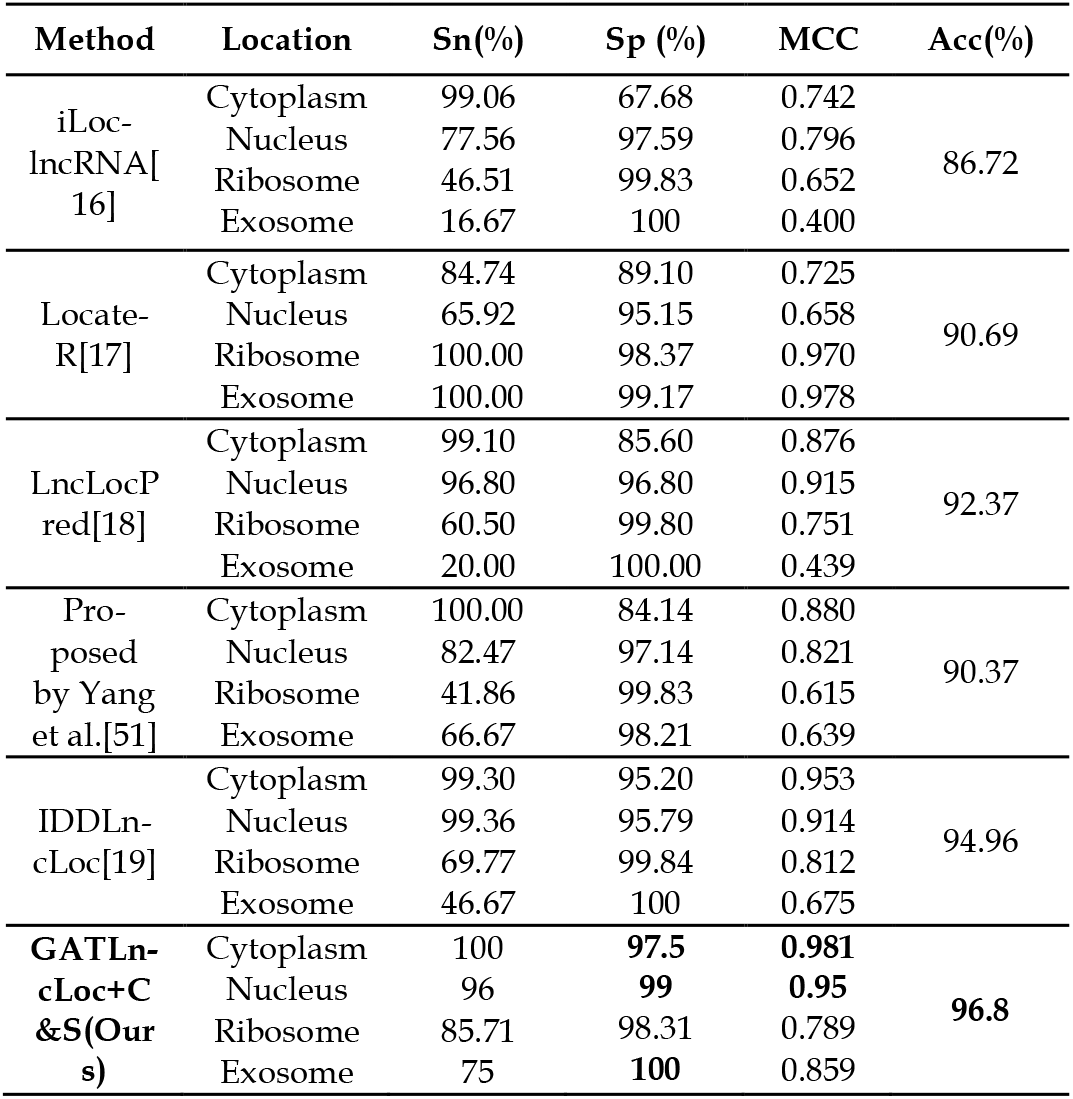
Comparison with state-of-the-art methods on 4 subcellular localization dataset.

There are three reasons that GATLncLoc+C&S outperforms previous models: first, GATLncLoc+C&S is based on a graph neural network, which can extract advanced features from the optimal characteristics of LncRNA sequences to complete the classification task, while most existing methods extract multifaceted features from LncRNA sequences and select the optimal features by achieving classification; second, we found the importance of the proportion of samples with the same localization label between neighborhood nodes, as this ratio increases, GATLncLoc+C&S can better learn the information of different LncRNAs in the same class of samples; finally, we proposed to use soft label correction graph structure to make more connections between representatives of the same localization label and better propagate label information.

### 3.8 New Node Prediction Experiment

For the prediction experiments based on new samples, we used the independent test set collected from the lncSLdb database and DeepLncLoc [20], then obtained a dataset of 60 LncRNAs by removing the sequences containing the special symbols “N, R, S, Y” (Table 7). The LncRNAs in the new dataset did not appear in the model training. In addition, the new nodes were added to the graph using the similarity calculation method proposed in this paper, and the new nodes were predicted directly by the pre-trained GATLncLoc+C&S model with an accuracy of 40% in the classification experiments for 5 subcellular locations and 53.3% in the 4 subcellular localization. Meanwhile, the new datasets are available at lncLocator (available at http://www.csbio.sjtu.edu.cn/bioinf/lncLocator/), iLoc-lncRNA (available at http://lingroup.cn/server/iLoc-LncRNA/download.php) and DeepLncLoc (HTTP://bioinformatics.csu.edu.cn/DeepLncLoc/) for prediction in the website. From Fig.9, it can be seen that the prediction accuracy of GATLncLoc+C&S for new nodes is higher than that of lncLocator and iLoc-lncRNA by about 10% and 8.33%, and obtains the same accuracy of 53.3% as DeepLncLoc. However, the website developed by DeepLncLoc can predict only one LncRNA at a time, while GATLncLoc+C&S is able to add multiple LncRNAs to the graph for prediction. These experimental results suggest that GATLncLoc+C&S not only has better generalization ability but also performs better in time efficiency. In general, the existing models performed poorly indirectly predicting new LncRNAs, mainly because the small number of training samples limited the generalization ability of the models.

**Table 7.** Statistics of the new dataset.

| Location | Dataset3 |
| --- | --- |
| Cytoplasm | 19 |
| Nucleus | 18 |
| Cytosol | 8 |
| Ribosome | 8 |
| Exosome | 7 |
| **Total** | **60** |

**Fig. 9.**
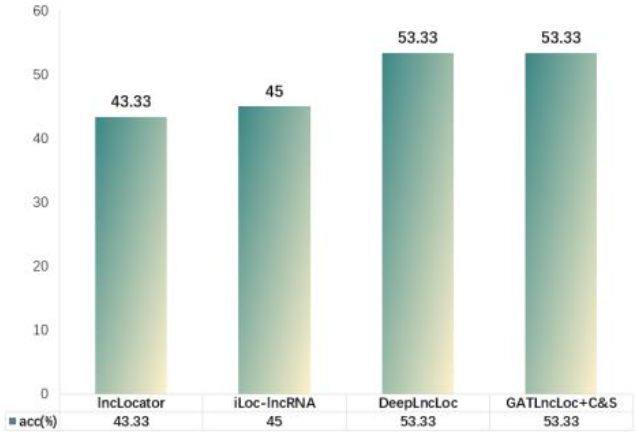
Comparison of prediction experiments for the new dataset.

## 4. Conclusion

Our proposed graph neural network and label propagation-based prediction model GATLncLoc+C&S is a novel approach to lncRNA subcellular localization. On the one hand, we consider the importance of the proportion of samples with the same localization label between neighborhood nodes before constructing the graph structure, which has not been paid attention to before; on the other hand, before propagating the label information, we propose that using soft label correction graph structure can correct the label more effectively during propagation and improve the prediction accuracy, and the model can adapt quickly to new samples. From the experimental results, GATLncLoc+C&S has a certain degree of contribution and competitiveness in solving LncRNA subcellular localization. In the subsequent study, we will think more about LncRNA subcellular multi-localization prediction.

**Xi Deng** is currently working toward the master’s degree in the School of Information, Yunnan Normal University, Kunming, China. Her current research interests include computational biology and machine learning.

**Lin Tang** is currently teaching at Yunnan Normal University, Kunming, China. His current research interests include machine learning, deep learning, and bioinformatics.

**Lin Liu** is currently teaching at Yunnan Normal University, Kunming, China. Her current research interests include machine learning, deep learning, and bioinformatics.

